# Circuit-Level Dynamics of Slow Wave Activity and Propagation During the Awakening Process

**DOI:** 10.1101/2024.12.20.629448

**Authors:** Antonio Pazienti, Mariel Müller, Conrado A. Bosman, Umberto Olcese, Maurizio Mattia

## Abstract

Slow-wave activity (SWA) is a hallmark of the loss of consciousness in non-REM sleep and anesthesia. The mechanistic underpinnings of SWA, and its evolution when transitioning towards the conscious brain state is poorly understood. We address this topic by recording multi-area and laminar activity in posterior parietal (PPC) and primary visual (V1) cortices of mice spontaneously awakening from isoflurane anesthesia. Spectral power is stronger in PPC (especially in superficial layers) during deep unconsciousness, but stronger in V1 when awakening. Rostro-caudal (feedback-like) propagation of SWA also shows state-dependent modulation, particularly in layer 5. The excitability of layer 2/3 neurons, hindered at high isoflurane, recovers during awakening, when V1 and the feedforward pathway reacquire a strong role. Detailing the hierarchical and laminar properties of spontaneous traveling oscillations, we provide evidence that SWA is a multiscale phenomenon. Explicating the functional role of these processes is critical to understand the neuronal mechanisms of consciousness.

## Introduction

Neuronal activity in brain networks unfolds across a multitude of spatiotemporal scales (Buzsaki, 2006). The large cortical network in particular shows a high flexibility in how information propagates. This process is mainly shaped by the brain hierarchical organization, but is also modulated by several factors, owing to the need to bidirectionally interact with the environment in a manner that also depends on the brain internal state. The bottom-up flow of information from sensory to associative and frontal cortices and the top-down feedback in the opposite direction follow specific and almost complementary laminar and inter-areal trajectories (Markov et al., 2013, 2014). The effectiveness in the transmission and transformation of information across these pathways underlies our capability of conscious processing (Koch et al., 2016a) and as such it is strongly affected by brain state (Massimini et al., 2009; Lewis et al., 2012; Bettinardi et al., 2015; Ishizawa et al., 2016). During wakefulness, recurrent interactions between cortical areas are prominent (Gilbert and Li, 2013; Koch et al., 2016b; Pennartz et al., 2023) and have been shown to be necessary for sensory-motor transformations to take place (Lamme and Roelfsema, 2000; Gilbert and Li, 2013; Pennartz et al., 2019). Such activity reverberation has also been hypothesized to be at the heart of conscious processing (Koch et al., 2016a; Storm et al., 2017; Olcese et al., 2018b). During brain states in which consciousness is lost - such as sleep and anesthesia - interactions between brain regions are generally dampened (Massimini et al., 2005; Sarasso et al., 2015; Olcese et al., 2016, 2018a), impairing the effective processing of sensory information (Sela et al., 2020; Hayat et al., 2022).

However, the mechanistic underpinnings of natural as well as externally-imposed alternations between conscious and unconscious states (e.g. the sleep/wake cycle and the induction of anesthesia, respectively) are still widely unknown. A key role is played by the thalamocortical system, which is modulated by but also determines the brain state (Steriade et al., 1993). Long-range functional connectivity between cortical areas is also widely modulated across global transitions in brain state, while local connectivity appears to be preserved (Olcese et al., 2016; Storm et al., 2017). Especially during non-REM (NREM) sleep – a brain state with characteristics very similar to those obtained with non-dissociative anesthetics (Mashour, 2024) – feedforward propagation of sensory information is more limited compared to wakefulness (Sela et al., 2020). However, waves of slow oscillatory dynamics (slow waves, which are the hallmark form of cortical activity during NREM sleep) typically originate in frontal cortical areas and propagate towards the back of the brain (Massimini et al., 2004). Slow waves have been linked to key functions associated to sleep such as memory consolidation (Huber et al., 2004; Marshall et al., 2006; Chauvette et al., 2012) and synaptic homeostasis (Tononi and Cirelli, 2014), and their properties and dynamics have been shown to vary as a function of the depth of sleep and anesthesia (Vyazovskiy et al., 2009). In the past decade, slow waves have also been shown to occur also during quiet wakefulness, either as local episodes of sleep (Nir et al., 2011; Vyazovskiy et al., 2011) or as a possible default state for cortical function in the absence of external and internal modulations (Sanchez-Vives and Mattia, 2014). This contrasts the mainstream view that the occurrence of slow waves represents a key marker of the loss of consciousness (Koch et al., 2016a). Also, what function slow waves might play outside of sleep is not fully understood (Vyazovskiy et al., 2011; Bhattacharya et al., 2022a). Finally, it remains unclear whether slow waves taking place during wakefulness intrinsically differ from those occurring during sleep and anesthesia (Andrillon et al., 2021), and how global state transitions such as between wakefulness and anesthesia unfold (Sulaman et al., 2023). Addressing all these questions is essential to uncover the architecture and function of one of the most fundamental forms of cortical activity underlying information propagation across brain states (Sanchez-Vives et al., 2017).

Here, we focused on the laminar spatiotemporal unfolding of isoflurane-induced slow-wave activity (SWA) and on its modifications during the awakening process in head-fixed mice. To this purpose, we simultaneously recorded the activity of several neuronal assemblies probed across all cortical layers in both a primary and a higher sensory cortex during the transitions from isoflurane-induced SWA to wakefulness. Our results show that specific frequency-related signatures characterize the restoration of consciousness from anesthesia. In particular, we observed a shift of power in the delta band. Delta activity was dominant in the posterior parietal cortex (PPC) at deep isoflurane levels (in particular for layer 5). In contrast, a shift to higher frequencies was observed towards awakening, when infragranular layers and primary visual cortex (V1) showed the largest activity amplitude. Furthermore, we found that the two main propagation modes of slow waves – from frontal to caudal areas and vice versa – differ in terms of microcircuit architecture, suggesting that they reflect distinct processes. Specifically, the firing rate of PPC layer 5 neurons and the involvement of infragranular neurons in feedback-propagating slow wave appear strongly modulated by the anesthesia level, while the feedforward propagation mode appears more stable. Overall, our results shed light on the characteristics of spontaneous oscillations in the transition from an unconscious to a conscious state, contributing to set a reference on future works on the hierarchical organization of the intact brain.

## Results

### Slow-wave propagation across cortical areas and layers

After performing craniotomy surgery and probe implantation (Fig. 1A, see Methods), we varied the level of isoflurane in a controlled way in 8 mice along two anesthetic concentrations (0.5% and 0.3%). Each concentration was maintained for approximately 30 minutes. Then, after setting isoflurane concentration to 0%, we let the mice awake naturally (Fig. 1A, bottom). For each concentration level, we discarded the first 15 minutes, as we assumed them to be a nonstationary response to the anesthesia change, and we analyzed the electrophysiological activity of the following 15 minutes only. Our electrodes probed all cortical layers in both primary visual cortex (V1) and posterior parietal cortex (PPC). From the electrophysiological signals recorded in the 64 channels (16 per shank, 2 shanks per area, 2 cortical areas) we extracted the local field potential (LFP, raw signal low-pass filtered at 200 Hz) and the multi-unit activity (MUA, computed here as the relative power of the detrended signal in the high-frequency domain [0.2, 1.5] kHz - see Material and Methods) (Fig. 1B). A clear alternation emerged between quiescent and active states (or Down and Up states, with relatively low and high rates of spike firing), underpinning the so-called slow-wave activity (SWA, Fig. 1B-C).

**Figure 1.**
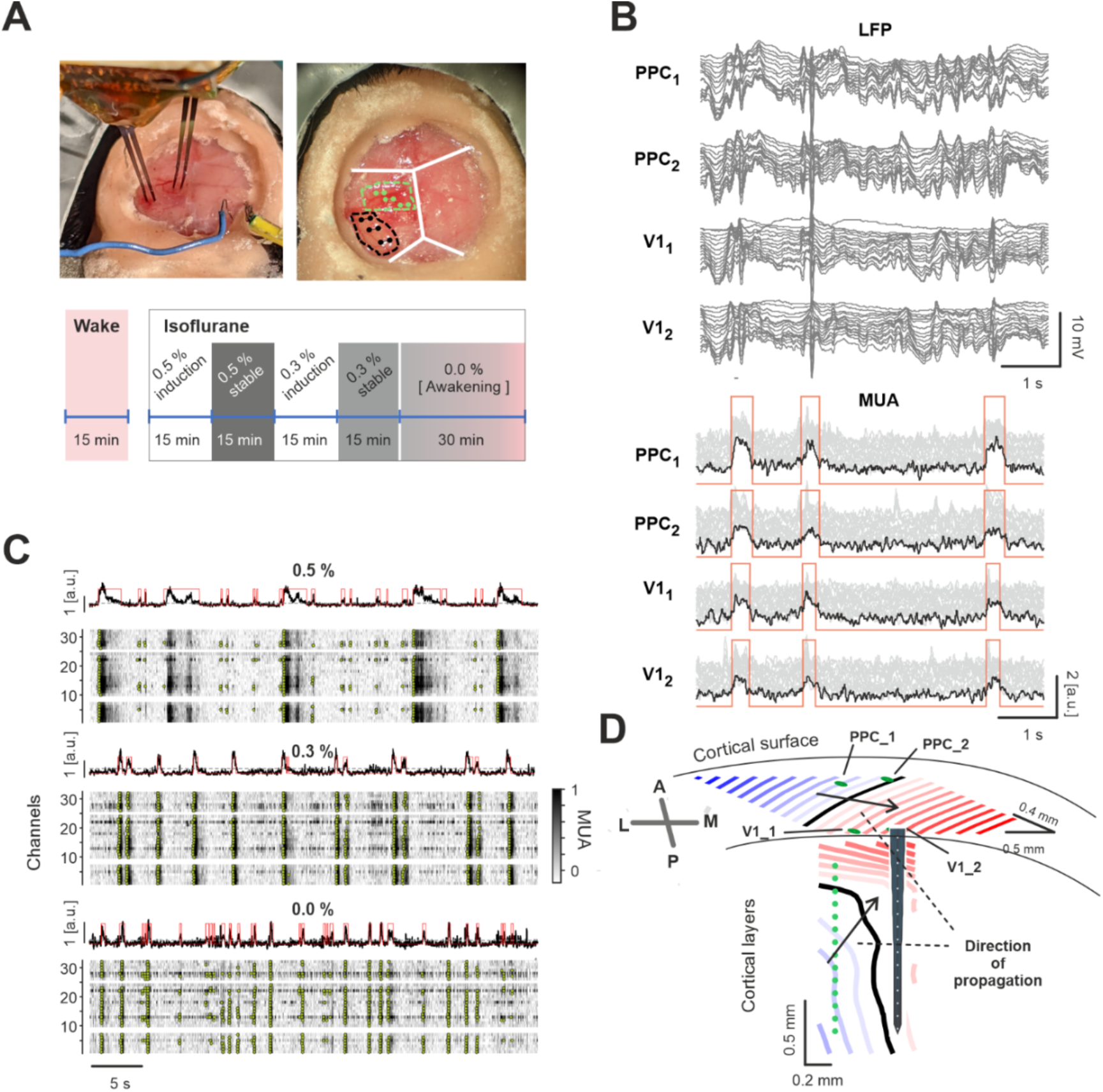
Electrophysiology and recordings. **A.** (Top) Electrode placement during a single session (left) and across multiple sessions (right). V1 insertions are marked in black, PPC insertions in green. White lines indicate the approximate location of skull sutures. (Bottom) Schematic of the anesthesia protocol. **B.** (Top) Representative low-frequency signal (LFP) for the different electrodes (groups of traces) and channels (traces within each group, ordered by channel depth on the electrode); (bottom) representative MUA traces (grey) and average value for each of the four electrodes (black). Red: identified Up states. Traces were shifted vertically for better visibility. **C.** Representative detection of Down-Up transitions for different isoflurane levels. From top to bottom: average firing rate (in arbitrary units) across a pair of electrodes (black) with Down-Up detections (red line); relative MUA of the 32 channels with detected Down-Up transition channel-wise (dots). **D.** Schematic representation of wavefront of slow-wave activity propagating (from blue to red) between cortical areas and among layers within a cortical column.

Using a previously developed approach, we detected the Down-to-Up transitions of MUA in each channel (Mattia et al., 2021; Tort-Colet et al., 2021; Pazienti et al., 2022) and computed their relative time lag, eventually grouping transitions estimated to belong to the same propagating wave as wavefronts (see Material and Methods). Individual wavefronts were then clustered based on the similarity of the channels’ activation (see Material and Methods). We measured how spiking activity propagated both between cortical areas and across the layers (Fig. 1D). We refer to “three-dimensional” propagation patterns as measuring the temporal lags of channel activations in the layers we obtained “vertical” flow (from deeper to more superficial layers or vice versa, Fig. 1D, bottom) and computing lags between the V1 and the PPC shanks we measured the “horizontal” propagation (Fig. 1D, top).

### Global state shapes SWA features

The aim of our study was the characterization of the organization of the SWA traveling across the cortical volume (layers and areas). According to previous studies, SWA originates in layer 5 (L5) and then propagates both to supragranular layers (Chauvette et al., 2010; Beltramo et al., 2013; Senzai et al., 2019; Mattia et al., 2021) and to other cortical areas (Mohajerani et al., 2013; Dasilva et al., 2021; Pazienti et al., 2022). Here we took a step further by detailing the relative timing and features of this complex, three-dimensional propagation.

We first focused our analysis on the characterization of SWA propagation across cortical areas (V1 to PPC and vice versa). We refer to this as cortico-cortical (or horizontal) propagation. To this purpose, we identified the electrode channels situated in L5 (see Materials and Methods) and extracted various relevant wave features (namely the transmission delay, the inter-wave interval [IWI] and the Up state duration) based on the Down-to-Up transitions measured in L5 (Fig. 2A). We quantified these features as a function of the global state induced by the anesthesia level. We will refer to the state associated with the highest concentration of isoflurane inducing SWA as “Mid” anesthesia, and we similarly termed “Light” anesthesia and “Awakening” the states associated with the steps of anesthetic concentration progressively fading out. For most of the experiments (8 out of 11) “Mid” corresponds to 0.5 % isoflurane, “Light” to 0.3 % and “Awakening” to 0 %. Three experiments were re-classified because they clearly showed features of another state (see Material and Methods for details). Consistently with previous work (Dasilva et al., 2021; Pazienti et al., 2022), the speed of intra-cortical propagation monotonically increased from about 10 mm/s to 30 mm/s as the anesthetic vanished (Fig. 2B). The Up state duration and the coefficient of variation of the inter-wave interval were also well correlated with the global brain state, decreasing and increasing, respectively (Fig. 2C,D). The frequency rate of the slow waves increased from 0.22 ± 0.05 𝐻𝑧 under Mid anesthesia to 0.39 ± 0.14 𝐻𝑧 in Light anesthesia (𝑝 < 0.05, Wilcoxon rank sum test). In the awakening phase the frequency of the Up states underwent large fluctuations, also due to the natural, progressive vanishing of the SWA. The frequency in this phase was 0.38 ± 0.20 𝐻𝑧 in the first third of the awakening phase and 0.23 ± 0.16 𝐻𝑧 in the last third.

**Figure 2.**
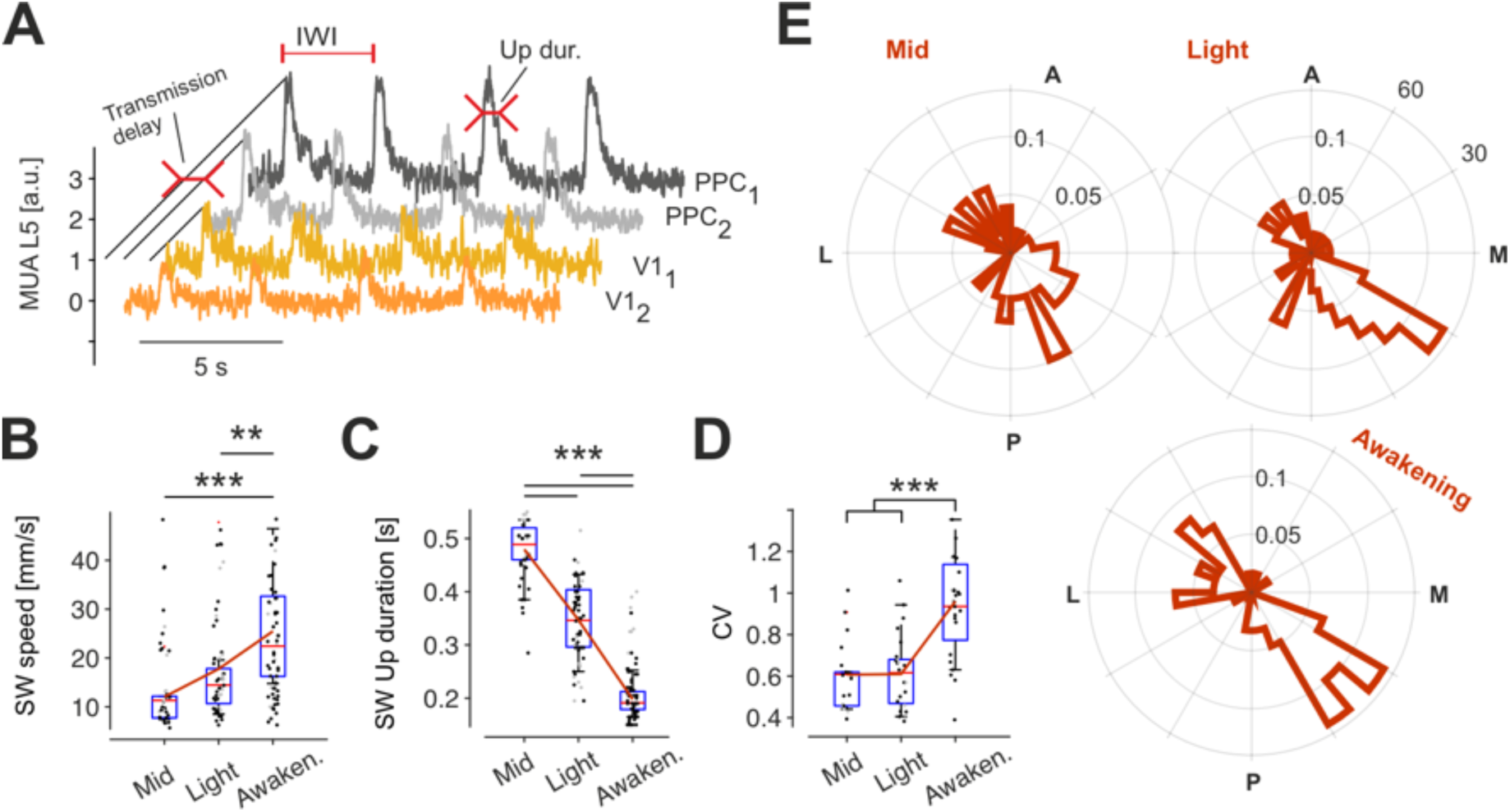
Slow-wave activity characterization. **A.** Representative MUA activity in L5 in the four recording cortical locations and description of the extraction of relevant features of the SWA (defined in panel B). **B-D.** Speed of cortical propagation (B), Up state duration (C) and coefficient of variation of the inter-wave interval (IWI) (D) as a function of anesthesia level. See panel A for the procedure of extraction of the features. **E.** Histograms of the probability distribution of the horizontal (i.e., cortico- cortical) directions of propagation for the three anesthesia levels.

We then investigated the direction of the wavefront propagation. In our data, consistently with previous observations (Dasilva et al., 2021; Pazienti et al., 2022), we observed two main directions, that is, antero-lateral to posterior-medial and vice versa (Fig. 2E). Interestingly, these main directions of propagations – back-to-front (feedforward) and front-to-back (feedback) – correspond to activity travelling from V1 to PPC and *vice versa*, usually associated to feedforward and feedback information flows, respectively.

### The laminar distribution of oscillatory dynamics rearranges across parietal and visual cortices when awakening from isoflurane

Having characterized horizontal slow-wave propagation along the awakening process, we aimed at investigating its properties across cortical layers. The contribution of specific frequency bands is long known to play a role in characterizing the transition from the unconscious to the conscious brain state (Akeju and Brown, 2017; Adamantidis et al., 2019). Induction and changes in the isoflurane concentration were clearly visible along our experimental protocol in terms of power spectral density (Fig. 3A). SWA-related spectral activity (normalized to the spectrum during wakefulness) centered around 0.2 Hz in the deepest anesthesia state (“Mid”, blue in Fig. 3B), that gradually switched to ∼1 Hz in Light anesthesia and finally to the theta band (∼6 Hz) in the awakening (Fig. 3B). This was quantified by the area of the spectrum in different frequency bands. The “infra-slow” band (0.06-0.25 Hz), observed also in ketamine-based anesthesia, see (Vanhatalo et al., 2004; Tort-Colet et al., 2021) dominated in the Mid state. The “slow” band (0.55-0.95 Hz) reached its maximum power in the Light state. The “theta” band (1-6 Hz) was reduced compared to awake in the Mid and Light states and re-emerged when anesthesia faded out (Fig. 3C). Interestingly, when comparing the power in the different cortical areas, we found that the power in the slow frequencies (sub-Hz, <1 Hz) was significantly stronger in PPC than in V1 (Fig. 3D, p<0.01).

**Figure 3.**
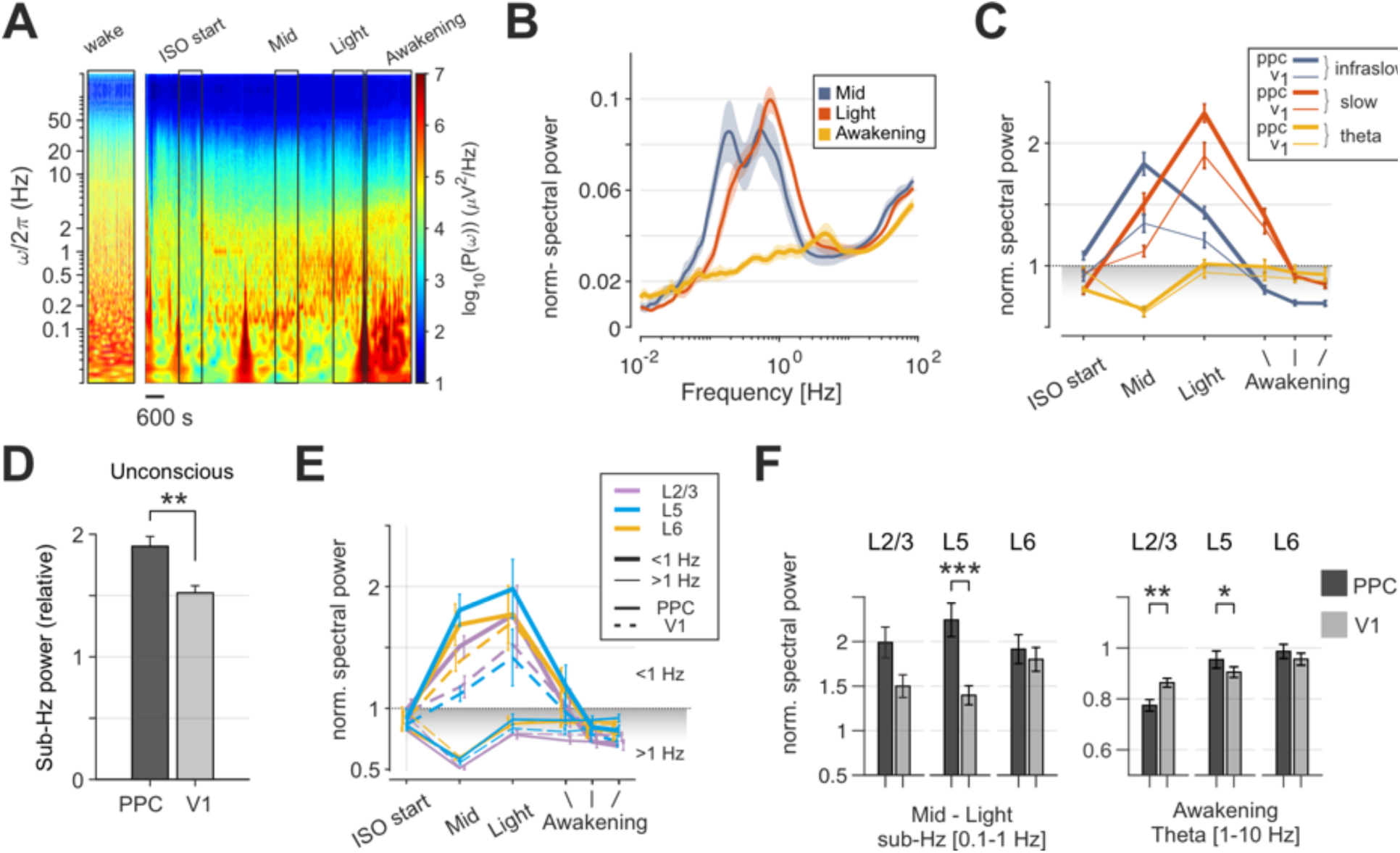
Spectral information in cortical areas and layers across brain states. **A.** Representative time-resolved power spectrum of the unfiltered local field potential along one recording session. Left panel, power during wakefulness, which is subsequently used as the reference for normalization. Right, black rectangles show periods of stable isoflurane concentration used for the analysis. Before the signal stabilized, burst of spectral activity involving slow frequencies are visible, presumably due to small movements of the animals. **B.** Example of power spectrum for one animal in the three global states, normalized for the power during wake. **C.** Total power of individual frequency bands as a function of the global state for both PPC and V1 areas. Infraslow: 0.13-0.25 Hz; slow: 0.55-0.95 Hz; theta: 1-6 Hz. **D.** Total power in the band 0.13-0.95 Hz band (referred to as sub-Hz) for PPC vs. V1. **E.** Total power in the sub-Hz band (thick lines, mostly above 1) and supra-Hz band (1-60 Hz, thin lines) for PPC (straight line) and V1 (dashed line) and all layers. **F.** Total power divided by layer for PPC (dark grey) and V1 (light grey) for sub-Hz bands in unconsciousness (left) and in the Theta band in awakening (right).

We then analyzed the contribution of the different layers, by identifying L2/3, L5 and L6 (see Methods). We observed a clear difference in the laminar distribution of the power. For all layers the sub-Hz power was stronger than power in the other bands in Mid and Light anesthesia, and the power >1 Hz was weaker than in the awake state, before eventually converging during awakening (Fig. 3E). Importantly, whereas PPC power dominated in deep anesthesia in the slow frequency bands (Fig. 3F, left), this was reversed in awakening, at least for supragranular layers (L2/3), for which V1 showed stronger power in the theta frequency band (Fig. 3F, right).

These results detail the laminar distribution of oscillatory power across frequency band in the mouse primary visual and parietal cortex, and demonstrate that different frequency bands are most prominent in either PPC or V1 depending on the global state.

### Feedforward and feedback SW propagation differently entrain cortical layers

Having determined the principal directions of propagation of the SWA between cortical areas, and the related changes in oscillatory dynamics occurring during the awakening process, we aimed at investigating more thoroughly the spatial organization of the SWA propagation across layers (intra-cortical laminar propagation) and between cortical areas (inter-areal horizontal propagation), and specifically whether these differed between the feed-forward and feedback SWA propagation modes.

For each of the two main directions of inter-areal SWA propagation (Fig. 4A), we tracked the three-dimensional propagation by taking as a reference the passage of the wavefront at layer 5 (Fig. 4B), where SWA is known to originate before spreading to the other cortical layers (Senzai et al., 2019; Mattia et al., 2021). For the feedback mode of propagation, the neuronal activity initiated in (at least one of) the PPC electrodes (Fig. 4, top row), whereas V1 activated earlier in the feedforward mode (Fig. 4, bottom row).

**Figure 4.**
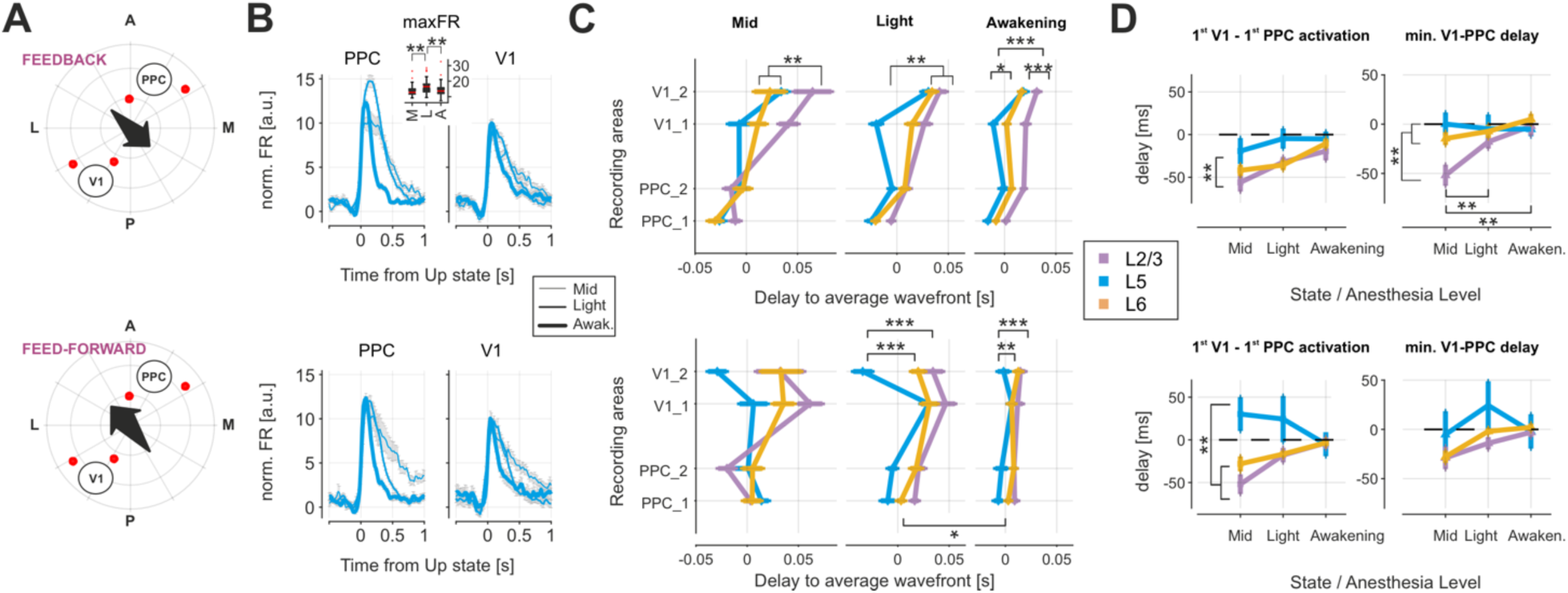
Laminar characteristics of slow-wave propagation across brain states. **A.** Cortico-cortical directions of slow-wave propagation correspond to top-down (feedback, top row) and bottom-up (feedforward, bottom row) processing. Red dots, positions of the 16-channels electrodes. **B.** Average Up state waveforms in L5 for all areas, consciousness states, and direction of propagations. **C.** Relative temporal delay of slow-wave propagation for each recorded cortical area for L5 (cyan), L2/3 (purple) and L6 (yellow) for the three global states. **D.** Relative temporal delay between the first activation of PPC and V1 areas, respectively (left), and of the minimal delay between PPC and V1 (right) as a function of the global state. The top row shows antero-lateral to posterior-medial (feedback) propagation; the bottom row shows posterior-medial to antero-lateral (feedforward) propagation.

Consistently with the results of the spectral power content (previous Section and Fig. 3) the activation of L5 in the SWA was stronger in PPC than V1 (Fig. 4B), and significantly modulated by the different global states. Interestingly, for feedforward propagation the firing rate in L5 did not undergo strong state-dependent modulations (Fig. 4B, bottom), whereas for the feedback mode the L5 maximal firing rate in PPC showed a reversed “U-shaped” modulation, being weak in the Mid state, maximal in the Light state, and intermediate in the Awakening state (Fig. 4B, top, inset).

Overall, while L5 activates before L2/3 and L6 in all states and for both modes of propagation (Fig. 4C), we noticed a significant difference in the involvement of the supra- and the infra-granular layers, with L2/3 being excited at a significant later point in time for all conditions but the Mid state of the feedforward mode (Fig. 4C, cf. the delay values of Fig. 4C bottom, Mid column). Furthermore, L6 activated almost synchronously to L5 for deeper isoflurane levels but accumulated a lag when the global state changed towards the awakening (cf. second and third columns of Fig. 4C, cyan vs. yellow lines).

We then investigated the time it took for the propagating spiking activity to travel through the cortex, by measuring the time delay from the first activation in V1 to the first activation in PPC and the minimum PPC- V1 delay irrespective of the order of activation (Fig. 4D). Here, negative delays mean that PPC activates before V1. Surprisingly, whereas in L5 the feedback and the feedforward modes of propagation differentiated themselves based on the order in which activity arose in PPC and V1, for both L2/3 and L6 activity always initiated in PPC. This is compatible with recent results showing that feedback projections from parietal cortex to V1 preferentially supragranular and L5 neurons (Aggarwal et al., 2022). In other words, synaptic transmission restricted to infragranular (supragranular) layers connecting primary visual and parietal cortical areas seems to be unaffected by mode of propagation. In all conditions, the delays rapidly decayed as a function of the global state (Fig. 4D, from Mid to Awakening), manifesting once again the increase of propagation speed as the brain transitions to consciousness. Furthermore, the quantification of the minimum time it takes for layer activation to travel from PPC to V1 (or *vice versa*) showed that, as this lag progressively decayed, for the feedback mode only layer 2/3 experienced a significant modulation when the awake state was approached (Fig. 4D; cf. top and bottom rows). That is, in the deeper unconscious level the L2/3 seems to be more severely affected than infragranular layers.

Our results show that in the feedback cortical propagation of slow waves the stronger effect of isoflurane-induced unconsciousness is a significant dose-dependent modulation of the Up-state in L5 neurons and of L2/3 excitability, whereas the latter rapidly recover during awakening, when V1 and the feedforward pathway reacquire a strong role.

## Discussion

The process of awakening from an unconscious brain state – whether natural sleep or drug-induced anesthesia – is associated with the transition from a relatively “simple” synchronized activity towards a more complex desynchronized state (Sanchez-Vives et al., 2017; Demertzi et al., 2019; Stevner et al., 2019). The rise of complex dynamics is thought to underlie the ability of the brain to process a complex environment, that requires it to produce context-dependent responses to the continuous barrage of sensory stimuli it receives (Sporns, 2022; Storm et al., 2024). The activity of the conscious brain can indeed be conceptualized as a synergistic interplay between self-generated and externally-driven information processing. These two modes of “computation” are associated to a dual counterstream organization. Feedforward (FF) and feedback (FB) pathways differently involve brain areas and, locally, cortical layers in in a hierarchically structured organization (Coogan and Burkhalter, 1990; Douglas and Martin, 2004; Markov et al., 2013; Vezoli et al., 2021; Aggarwal et al., 2022; Barzegaran and Plomp, 2022; Spyropoulos et al., 2024).

Here we reported the first detailed description of the three-dimensional propagation of slow-wave activity, that is involving at the same time two relevant stations of the cortical sensorial pathway (V1 and PPC) and their local circuitry — cortical layers. Slow waves use the FF and FB propagation modes to engage the cortical layers in a way that depends on the level of unconsciousness. Notable differences between the present study and previous works on cortical propagation of activity are i) the use of the unconscious (rather than the awake) state, and ii) the focus on spontaneous FF/FB propagation, unrelated to external stimuli or stimulation as in previous studies (see for instance Aggarwal et al., 2022; Barzegaran and Plomp, 2022; Shen et al., 2022; Spyropoulos et al., 2024).

Feedforward propagation in the neocortex is known to reflect the processing of sensory stimuli, is mediated by thalamic innervation (Douglas and Martin, 2004; Balcioglu et al., 2023). Feedback signals are instead known to travel both superficially (supragranular to supragranular layers) and in deep layers (infragranular to infragranular layers), and they are believed to modulate the sensory pathway (affecting the FF-traveling signals) (Aggarwal et al., 2022; Barzegaran and Plomp, 2022). We have shown that slow wave activity similarly follows a feedforward and a feedback pathway, from V1 to PPC and vice versa. During feedback propagation we document a modulation of the L5 firing rates as a function of anesthesia depth (Fig. 4B). This suggests that feedback pathways are more sensitive to the brain’s overall state (Hudetz et al., 2020; Suzuki and Larkum, 2020), and is consistent with the pivotal role played by L5 neurons in mice under anesthesia (Bharioke et al., 2022). In contrast, the activation of L5 in the FF mode does not show any modulation based on global states. This relative stability indicates that the forward pathway may provide a more consistent driving force for neuronal activity in L5 compared to the dynamic changes observed in the complementary pathways. As for supragranular (SG) layers, we observed that these experience the stronger effect of isoflurane in FB propagation (Fig. 4D, top), whereas the transmission is much hindered at deep isoflurane levels. Both for FB and FF propagation, supragranular and infragranular (IG) layers activate first in PPC (Fig. 4D), irrespective of the direction of propagation. This is consistent with the fact that the L5 network synchronizes independently of the other layers and might be due to the enhanced laminar propagation reported in resting states as compared to wake (Kharas et al., 2022).

A signature of sleep and anesthetized states is the presence of network activity in the slow frequency band, that is up to ∼5 Hz (Sanchez-Vives et al., 2017; Bhattacharya et al., 2022b). Consistently with this, we observed increased power in the infra-slow (∼0.2 Hz) and slow frequency bands under anesthesia. Our present results detail more thoroughly how these oscillations are modulated in the transition from unconsciousness to consciousness. In particular, the spectral power of these oscillations switched from slower to higher frequencies when the isoflurane concentration was lowered, and it was consistently higher in PPC than V1 during mid and light anesthesia. Importantly, we showed that the power in superficial layers is stronger in PPC in deeper unconsciousness states but stronger in V1 in awakening, coherently with the fact that SG layers mostly show feedforward projections.

As a consequence of our approach – investigating spontaneous propagation in the slow oscillation state – our results differentiate from what is known from the FF and FB pathways during perception and task execution. A key difference in the way cortical circuits organize their spatiotemporal activity in the unconscious brain compared to the feedforward and feedback information flow emerging in the awake brain relates to the propagation speed of the activation waves. Indeed, ranging from passive sensing via whisker deflection (Ferezou et al., 2007) to active visuo-motor integration (Peters et al., 2022), evoked activity propagates from sensory to motor cortices of behaving mice with a latency ranging from 15 ms to 30 ms, respectively for the somatosensory (Constantinople and Bruno, 2013) and visual (Kirchberger et al., 2023) system. By considering a cortical distance of about 5 mm between visual and motor areas of the same hemisphere (Xiao et al., 2021), activity is then expected to travel at the lowest at 5 mm / 30 ms ≃ 170 mm/s. This estimate appears to be an underestimate of recent experimental evidence reported for feedforward waves of gamma activity (30-50 Hz, tightly related to local multi-unit activity), which, in response to a visual stimulus, arise in primary visual cortex (V1) and propagate towards frontal area at a median velocity of 800 mm/s (Aggarwal et al., 2022). The slow waves of MUA activation that we characterized here and in previous studies propagate at a maximum mean velocity of about 30 mm/s (Ruiz-Mejias et al., 2011; Stroh et al., 2013; Shimaoka et al., 2017; Pazienti et al., 2022). Slow-waves are thus one-order of magnitude slower compared to the activity propagation observed in the feedforward mode during wakefulness. Our results then support the hypothesis that, although in the conscious and unconscious brain both feedforward and feedback modes of propagation naturally engage the cortical network, they are associated to a different functional organization.

Another important difference with respect to previous results lies in the cortical propagation of oscillatory activity. Under anesthesia this is sustained by a direct communication between nearby L5 assemblies (Fig. 4), for both the feedforward and the feedback modes. Activation of L6 and L2/3 follows L5 activation, irrespective of propagation direction. This aligns with the previously characterized relevance of L5 in the generation of Up states (Chauvette et al., 2010; Beltramo et al., 2013; Mattia et al., 2021; Tort-Colet et al., 2021; Bharioke et al., 2022). In addition, isoflurane was shown to differentiate from other anesthetics, in that it synchronizes also neurons in L1, L4 and L6 (Bharioke et al., 2022). This scenario and our results, however, contrast with what is known for activity propagation in the awake state. In behaving animals, as noted above, information flow is known to preferentially follow the supragranular-to-granular pathway for FF and the parallel infra- and supragranular pathways for FB (Douglas et al., 1989; Coogan and Burkhalter, 1990; Douglas and Martin, 2004; Markov et al., 2013; Hudetz et al., 2020). For example, feedforward inter-areal communication in rodents is thought to be primarily mediated by both L2/3 and L5 (Harris and Shepherd, 2015; Aggarwal et al., 2022), in particular, for what pertains communication between V1 and PPC (Glickfeld et al., 2013; Olcese et al., 2013; Yang et al., 2013). Feedback projections engage both L23 and L5/6, and different roles have been proposed for the different projections (Hudetz et al., 2020). All this is in stark contrast with the L5-mediated propagation that we observed here for traveling slow waves.

Our results suggest the emergence, over the course of awakening, of a more local computation of the cortico-cortical input received both by primary and higher cortical areas. This supports the view that a disruption in precise (locally-specific) information transfer, in particular affecting feedback slow wave propagation and with PPC appearing as a key hub, correlates with the loss of consciousness occurring in anesthesia. Indeed, we observed marked differences (dominance of L5 computation, and partial disruption of laminar processing) with the modes of activation typically associated with conscious processing, which question the validity of considering slow wave propagation during anesthesia as a direct counterpart of awake cortical dynamics. Overall, rostro-caudal propagation leads to a more complex and dynamic activation pattern characterized by state-dependent modulation, whereas the posterior-frontal travelling mode results in a more stable and consistent activation profile. This distinction highlights the different roles these pathways have been proposed to play in cortical processing and signal transmission.

Our results show that the propagation of both back-to-front and front-to-back propagating slow waves mainly impinges on L5. Over the course of the awakening from anesthesia the propagation of feedback waves undergoes variations that are mostly visible in PPC. While it would be intriguing to draw parallel to waves of activity propagating along analogous routes in the awake brain – and thus draw conclusions about the relevance for sustaining conscious processing, we are aware of the critical differences in the circuit-level mechanisms of activity propagation differentiating drug-induced unconsciousness and wakefulness. These differences suggest that the propagation of (endogenously generated) slow waves in anesthesia and (exogenously generated) sensory- and task-related activity in the awake state are fundamentally distinct processes. This bears great relevance for developing novel indicators of the level of consciousness, that cannot simply be based on what we know about conscious processing during wakefulness. However, a more comprehensive understanding of the awakening process would require focus on natural sleep besides anesthesia and perform recordings with a denser spatial and temporal sampling, possibly involving higher cortical and extending also to subcortical areas – such as what is now made possible by Neuropixels probes (Steinmetz et al., 2021) and similar technologies. We believe that this kind of approach would finally make it possible to go beyond the limitations of the existing indicators of consciousness.

## Acknowledgements

This study was supported by the European Union’s Horizon 2020 Framework Program for Research and Innovation under the Specific Grant Agreement 785907 (Human Brain Project SGA2) and 945539 (Human Brain Project SGA3) to CAB and UO, by an Amsterdam Neuroscience Proof of Concept grant to UO, by intramural funds from the Swammerdam Institute for Life Sciences of the University of Amsterdam to UO and CAB, and by the European Union – NextGenerationEU and the Ministry of University and Research (MUR), Italian National Recovery and Resilience Plan (PNRR), M4C2I1.3, project ‘MNESYS’ (PE00000006) to MM.

## Materials and Methods

### EXPERIMENTAL MODEL DETAILS

#### Subjects

All the experiments performed for this study were approved by the Dutch Commission for Animal Experiments (Centrale Commissie Dierproeven) and by the Animal Welfare Body of the University of Amsterdam, under permit AVD1110020172385. A total of 4 adult male mice were used for this experiment, each for 3 consecutive days. Mice were about three months old at the time of experiments. In order to reduce the number of animal used in experiments, the same transgenic mice Pvalb-IRES-Cre (B6;129P2- Pvalb<tm1(cre)Arbr>/J) mice (Mus musculus) were used both for this and for other, unrelated experiments. Mice underwent a surgery to mediate the expression of Channelrhodopsin2 in parvalbumin-positive interneurons. This procedure is not further described as it does not pertain the current manuscript. However, it is important to mention that this transgenic line is often used to study the neuronal mechanisms of sensory processing and decision making in both awake and anesthetized mice (Olcese et al., 2013; Oude Lohuis et al., 2022b, 2022a). Mice were kept on a 12 h reverse day-night cycle (lights off at 08:00hs and on at 20:00hs) with controlled values of humidity (55 % ±10) and room temperature (21.5 ± 2.0 deg). Food and water were provided ad libitum.

#### Surgical procedures

Following a week of habituation to handling mice were implanted with a light-weight titanium headbar aiming for head-fixation. All mice were anesthetized with Isoflurane (IsoFlo, 250ml; induction: 3-4%; maintenance: 1.5%). Carprofen (5mg/kg s.c.) was used as an analgesic. Breathing rhythm and rectal temperature - maintained at 37 deg - were monitored during the whole procedure. An ophthalmic ointment (Ophtosan, ASTfarma) was used for protecting the eyes. Hair was removed from the scalp with a hair removal cream (Veet), after which a local anaesthetic (Xylocaine 10%) and iodopovidone (antiseptic, Betadine, 30ml, Mylan) were applied to the skin. The scalp was then removed and the underlying skull was dried, softly scratched and covered by a layer of cyanoacrylate glue (Loctite 458). The head bar was attached to the skull via the use of dental acrylic (Superbond C&B) and dental cement (shade 528/1 pink, Kemdent). Following headbar implanation, the locations of the primary visual cortex V1 and of the posterior parietal cortex PPC were identified via the use of intrinsic optical signal imaging (IOI), as previously described (Olcese et al., 2013; Oude Lohuis et al., 2022b, 2022a). In a fraction of animals, a viral injection was done via the opening of a small craniotomy over V1 and through the injection of an AAV vector mediating the Cre-dependent expression of Channelrhodopsin (Oude Lohuis et al., 2022b, 2022a). This procedure was not part of the experiments presented here. The craniotomy and the skull were protected by a removable silicon layer (Kwik-Kast). After surgery, mice could recover for a week and were regularly monitored.

### METHOD DETAILS

#### Experimental procedure

Mice were first habituated to the experimental setup over the course of about two weeks, during which they were head-fixated for progressively longer periods, until they could stay in the setup without visible signs of stress for at least one hour. Once habituated, two small craniotomies were performed in correspondence of V1 and PPC, whose location was previously identified via IOI. Craniotomies were performed under isoflurane anesthesia, which was induced and maintained as described earlier. Animals were allowed to recover for one day. Two laminar silicon probes (Neuronexus A2x16-10mm-100-500-177) were then implanted in the two craniotomies. Each probe had two shanks, spanning a depth of 1500 um, with an inter-shank spacing of 500 um. Each shank had 16 recording sites, with an inter-site spacing of 100 um. The probes were acutely inserted to a depth of approximately 1500 um, in order to reach all cortical layers. The depth of the different recording sites was estimated via electrophysiological markers, described more in depth later. The relative position of the probes (Fig. 1A-B) was measured by aligning the micromanipulators used to position the probes to the same reference point on the skull (typically bregma). Probes were positioned such that the two shanks were oriented orthogonally to the caudo-ventral axis (see Fig. 1A). The location of probes in V1 and PPC was verified by coating the probes in DiI and by then aligning the insertion traces with a reference atlas through histological reconstruction (Fig. S1). Probe implantation was done under 1.5% isoflurane anesthesia, after which the actual experimental protocol started (Fig. 1A). First, isoflurane concentration was lowered to 0.5% and it was allowed to settle for 15 minutes. Recordings continued for 15 more minutes in a stable 0.5% condition. Isoflurane concentration was then lowered to 0.3%, and the same temporal procedure (2 x 15 minutes) was repeated. Finally, isoflurane concentration was reduced to 0% and the animals were allowed to wake up. Recordings continued for a further 30 minutes.

At the end of a recording session, probes were extracted, and the exposed cortex and skulls were again protected by a silicon layer (Kwik-Kast). The procedure was repeated over the course of at most 3 days, while slightly changing the location of probes (within each craniotomy) – see Fig. 1B. Probes were stained with DiI to enable post-mortem histological reconstruction.

#### Data acquisition

All data was acquired with a 64-channel Digital Lynx SX (Neuralynx) and digitized at 30 kHz, with a high pass filter set at 0.1 Hz.

#### Histology

At the end of experiments, animals were sacrificed by pentobarbital overdose and perfused with 4% PFA in phosphate buffer. Brains were sectioned and stained with DAPI.

### QUANTIFICATION AND STATISTICAL ANALYSIS

#### MUA estimate and detection of the state transitions

Up and Down states and multi-unit activity (MUA, also referred to as normalized FR) were estimated from the recorded raw signals (Supèr and Roelfsema, 2005; Stark and Abeles, 2007; Mattia et al., 2010; Reig et al., 2010; Sanchez-Vives et al., 2010). Briefly, the power spectra were computed from 5-ms sliding windows of the raw signal. MUA was estimated as the relative change of the power in the [0.2, 1.5] kHz frequency band. Such spectral estimate of the MUA was not affected by the electrode filtering properties, and it provided a good estimate of the relative firing rate of the pool of neurons nearby the electrode tip. In a second step, the MUA was logarithmically scaled to compensate for the high positive fluctuations due to the neurons closest to the electrode, thus obtaining the log(MUA) signal. We then smoothed the log(MUA) by performing a moving average with a sliding window of 40 𝑚𝑠. The average activity during Down states was subtracted from the resulting log(MUA) signal, so that the activity associated with the Down state fluctuates around 0. In the article, we refer to this signal as the normalized FR. From the long-tailed histogram of log(MUA), an optimal threshold separating Up and Down activity states was set at the absolute value of 0.4. The threshold for detecting both Down-to-Up and Up-to-Down transitions was the same.

The frequency of the slow waves was defined as the inverse of the duration of the entire Up-Down cycle. We reconstructed the activation waves from the detected Down-to-Up state transitions as in (Capone et al., 2019a) and in (Dasilva et al., 2021). Briefly, each wave was reconstructed pooling together the transitions occurring in multiple electrodes in a reasonable time interval that was iteratively reduced until each wave contained no more than one transition per channel. Each reconstructed wave was associated with a vector of relative time lags computed as the difference between the time of occurrence of the wave in each electrode and the average time of occurrence across all the electrodes taking part in the wave propagation. (This delay is referred to as transmission delay in Fig. 2A.) We rejected the waves occurring in less than twelve channels, otherwise we replaced the missing values with the result of a nearest-neighbor interpolation using the five nearest points in terms of Euclidian distance of wavefronts/time lag vectors. Using the resulting vectors describing the detected activation wavefronts, for each animal and anesthesia level we composed a time-lag matrix (TLM) having a number of rows equal to the number of waves and a number of columns equal to the number of recording channels of the experiment. We refer to the rows of the TLM, representing relative Down-Up transitions for each channel, also as time-lag arrays. The spatiotemporal course of the wavefronts was then obtained by spatially interpolating the time lags without smoothing, using a thin-plate spline method.

#### Modes of propagation and slow-wave features

Relying on the first principal component of TLM we clustered waves/rows sharing similar features with a k-means algorithm. The optimal number of clusters to be used to group the waves in each anesthesia level was chosen using a Silhouette method. For each cluster, we obtained the mean wavefront and its spatiotemporal profile by averaging the waves belonging to it. The mean wavefront extracted from each cluster was used to compute the mean velocity and direction of propagation of the cluster as in (Capone et al., 2019b). This was done relying on the smoothed surface T(x,y) (thin-plate smoothing spline) of the relative time lags associated with each cluster and distributed according to the electrode positions, and computing the local velocity as V(x,y)=1/(∂_x T^2+∂_y T^2 ). The gradient of this field points out the direction of the wave propagation. Mean values are computed across the 16 electrode positions. This procedure was used for example to produce the planes showing the progression of the wavefronts both interlaminar and tangential in Fig. 1D.

As for the other physiological quantities extracted from the slow waves, the maximal FR was defined for each Up state as the maximum of the FR during the Up subtracted by the baseline, defined as the average FR in the period between 200 and 50 ms before the transition to the Up state. Up duration was defined as the duration in seconds from the Down-Up and the Up-to-Down transition computed at half the baseline-corrected maximum FR. The inter-wave interval (IWI) was the time before two consecutive Down-Up transitions. The frequency of the slow wave was the inverse of this duration. The coefficient of variation was the ratio between the standard deviation of the IWI and their mean in a certain time segments.

#### Relative position of the recording sites

The precise relative position of the recording probes (and corresponding shanks) was determined by using photos of the recording setup taken on each recording session after the probes were inserted in the tissue. The image contained a measure reference (typically a piece of graph paper). The images were corrected for distortions offline using tools of the software CorelDRAW (version 22, Corel Corporation).

The distance between single channels in each shank (vertical distance) was fixed and given by the used probes (see above). This amounted to 100 𝑚𝑚.

#### Layer identification

In order to identify the channels associated to each of the relevant cortical layers we first identified the putative layer V from each of the four 16-channels shanks. L5 is known to have a leading role in the generation of the slow waves (Sanchez-Vives and McCormick, 2000). For each experiment session, using self-written code we therefore identified the channel activating first in the Down-to-Up transition and label it as L5. The labels obtained in this way were consistent with the method proposed by (Senzai et al., 2019). We used the latter reference and well-known further literature to assign channels to L2/3 and L6 on the basis of the distance from L5. This distance was 200 𝑚𝑚 for L2/3 and −100 𝑚𝑚 for L6, where positive and negative is towards the cortical surface and deeper, respectively.

#### Spectral analysis

Frequency analysis shown in Fig. 3 was performed using the continuous wavelet transform (CWT) in MATLAB (The MathWorks, Natick, MA). The CWT is obtained using the analytic Morse wavelet with the symmetry parameter, gamma, equal to 3 and the time-bandwidth product equal to 60. CWT uses 10 voices per octave. The minimum and maximum scales are determined automatically based on the energy spread of the wavelet in frequency and time. The power spectra in different conditions were normalized by dividing the wavelet decomposition for the corresponding decompositions in awake (i.e., before the isoflurane was inhaled).

#### Statistical analysis

All the analyses were performed in MATLAB (The MathWorks, Natick, MA).

